# Sleep spindle density and temporal clustering are associated with sleep-dependent memory consolidation in Parkinson’s disease

**DOI:** 10.1101/2023.10.20.563149

**Authors:** Soraya Lahlou, Marta Kaminska, Julien Doyon, Julie Carrier, Madeleine Sharp

## Abstract

Sleep is required for successful memory consolidation. Sleep spindles, bursts of oscillatory activity occurring during non-REM sleep, are known to be crucial for this process and, recently, it has been proposed that the temporal organization of spindles into clusters might additionally play a role in memory consolidation. In Parkinson’s disease, spindle activity is reduced, and this reduction has been found to be predictive of cognitive decline. However, it remains unknown whether alterations in sleep spindles in Parkinson’s disease are predictive of sleep-dependent cognitive processes like memory consolidation, leaving open questions about the possible mechanisms linking sleep and more general cognitive state in Parkinson’s patients. The current study sought to fill this gap by recording overnight polysomnography and measuring overnight declarative memory consolidation in a sample of thirty-five Parkinson’s patients. Memory consolidation was measured using a verbal paired-associates task administered before and after the night of recorded sleep. We found that lower sleep spindle density at frontal leads during non-REM stage 3 was associated with worse overnight declarative memory consolidation. We also found that patients who showed less temporal clustering of spindles exhibited worse declarative memory consolidation. These results suggest alterations to sleep spindles, which are known to be a consequence of Parkinson’s disease, might represent a mechanism by which poor sleep leads to worse cognitive function in Parkinson’s patients.

**Statement of significance:** Sleep — particularly spindle activity — is critical for memory consolidation, a core cognitive process. Changes to the architecture and oscillations of sleep are well documented in Parkinson’s disease (PD) and have been associated with worse overall cognition. However, whether altered sleep plays a causal role in this relationship, by directly interfering with sleep-dependent cognitive processes, or whether it represents a mere epiphenomenon of advancing disease, remains unknown. Our study is the first to investigate a possible direct relationship between sleep and cognition in PD. We show that sleep spindles and their temporal clustering into ‘trains’ relate to impairments in overnight declarative memory consolidation in patients. These findings are an important first step towards identifying modifiable sources of cognitive impairment in PD.

## Introduction

Disturbed sleep is common in Parkinson’s disease ^1–4^ and has been linked to cognitive dysfunction ^5–8^. Recently, there has been growing interest in examining alterations in sleep microarchitecture because specific neural oscillations of sleep have been proposed as potential biomarkers of cognitive state in Parkinson’s disease ^9,10^. However, the mechanisms linking sleep microarchitecture to cognition in Parkinson’s patients remain unclear. In particular, whether altered sleep plays a direct causal role in the cognitive deficits of Parkinson’s disease by interfering with sleep-dependent cognitive processes remains unknown.

It has repeatedly been shown that Parkinson’s patients have fewer sleep spindles, a type of fast oscillation with a frequency in the range of 11 to 16 Hz that occurs in non-rapid eye movement (NREM) stage 2 and stage 3 sleep ^11,12^. This observation stands out because sleep spindles are well-established to play a role in memory consolidation ^13,14^. Research in healthy adults has shown that greater spindle density (the number of spindles per minute of sleep) is associated with better sleep-dependent memory consolidation of both procedural ^15,16^ and declarative memories ^13,17–19^. Sleep spindles are thought to reflect bursts of thalamo-cortical neuronal activity, which in turn are thought to promote the neuronal plasticity necessary for memory consolidation ^20–22^. Additional evidence in support of the role of spindles in memory comes from studies in which sleep was pharmacologically manipulated with zolpidem. The increase in spindle count caused by zolpidem was associated with enhanced next-day memory performance in both healthy adults and individuals with schizophrenia ^23,24^. To date, no study has investigated the relationship between sleep microarchitecture changes that occur in Parkinson’s disease and specific sleep-dependent cognitive processes. Meanwhile, identifying such a link could provide a potential mechanism to explain the association observed in Parkinson’s disease between sleep microarchitecture and overall cognitive function.

The temporal dynamics and clustering of sleep spindles may also be important for memory consolidation. It has recently been shown that some spindles occur in quick succession, a phenomenon referred to as ‘spindle trains’ ^25^. This temporal clustering has been proposed as another mechanism of sleep-dependent memory consolidation because the repeated occurrence of spindles is believed to reflect repetitive reactivation of the memory engram, hence allowing for re-processing and further consolidation of memories ^25^. In young adults, the occurrence of spindles in trains has been associated with better overnight motor memory consolidation ^26,27^. In older adults, a similar relationship has been observed for declarative memory consolidation ^28^.

In the present study we aimed to determine if sleep spindles and their temporal dynamics are associated with overnight declarative memory consolidation in patients with Parkinson’s disease. We measured overnight sleep microarchitecture and overnight declarative memory consolidation using a standard verbal paired-associates memory task in a group of patients with Parkinson’s disease. We hypothesized that lower sleep spindle density and lesser clustering of spindles into trains during NREM sleep would be associated with worse overnight memory consolidation in Parkinson’s patients. Our results were broadly consistent with these hypotheses but the relationships we observed between sleep spindles and declarative memory consolidation were specific to NREM-3. By demonstrating a link between sleep micro-architecture and performance on a sleep-dependent cognitive process in Parkinson’s disease, our results raise the possibility that impaired sleep directly contributes to overall poor cognitive function in Parkinson’s patients by interfering with specific sleep-dependent cognitive processes.

## Materials and methods

### Participants

Forty-five patients with Parkinson’s disease were recruited from the McGill University Hospital Centre and from the Quebec Parkinson Network, a registry of patients interested in research. All participants had a diagnosis of Parkinson’s disease confirmed by a neurologist and were taking dopaminergic medications for their PD symptoms. Importantly, none were taking overnight dopaminergic medications. Participants were recruited as part of a larger multi-night interventional study on the effects of overnight levodopa on obstructive sleep apnea, the results of which will be published separately. Participants in the present study were tested only on the initial overnight screening visit of that study (i.e., before the initiation of the intervention), which served the purpose of identifying patients with obstructive sleep apnea for enrolment in the interventional study. Because our study took place on the screening visit, our sample included patients with and without obstructive sleep apnea (65% with moderate or severe OSA), which is consistent with previously reported estimates of OSA in Parkinson’s disease ^29^. None had major health issues, neurological disorders other than Parkinson’s disease or active psychiatric disorders. Six participants had missing memory recall data and two had an EEG recording of bad quality based on visual inspection. Thirty-seven participants were thus retained for the analyses. Two participants were excluded because they were identified as outliers based on their memory performance before sleep (recalled 0 and 3 out of 25 words). The final sample consisted of thirty-five participants. Demographic and clinical characteristics of the final sample are presented in Table 1. All participants provided written consent and were compensated for their participation. The study was approved by the McGill University Health Centre Research Ethics Board and all procedures were performed in accordance with the appropriate institutional guidelines.

**Table 1.** Demographic and clinical characteristics.

|  | Mean (range) |
| --- | --- |
| Age | 65.5 (47 - 81) |
| Sex, % men | 65.7% |
| Education (years) | 15.8 (9 - 22) |
| MoCA<sup>a</sup> | 26 (18 - 30) |
| Digit span (total)<sup>b</sup> | 11.5 (8 - 16) |
| UPDRS-III<sup>c</sup> | 23.3 (7 – 51) |
| Disease duration | 3.4 (0 – 26) |
| LEED<sup>d</sup>, mg | 492.7 (200 – 11,641) |
| Apnea-Hypopnea index | 23.6 (2.3 – 70.5) |

Table shows mean with the range in parantheses.
<sup>a</sup> MoCA = Montreal Cognitive Assessment
<sup>b</sup> Digit span total represents the sum of forward and backward digit span
<sup>c</sup> UPDRS-III = Unified Parkinson's Disease Rating Scale Part III, tested ON dopamine medication
<sup>d</sup> Levodopa equivalent dosing; includes levodopa, dopamine agonists, amantadine, monoamine oxidase inhibitors and catechol-O-methyl transferase inhibitors

### Overall procedure

All participants completed questionnaires about their clinical history and underwent a Unified Parkinson Disease Rating Scale – Part III motor assessment ^30^ and a Montreal Cognitive Assessment (MoCA; ^31^) during the evening portion of their overnight visit. As detailed below, memory testing took place in the evening and the following morning. Patients were instructed to take their usual dopaminergic medications 45-60 minutes prior to both sessions in order to control for the effects of dopamine medication state on recall. The evening session was scheduled between 5 and 6 PM to ensure that the dopaminergic medications were taken no later than 5 PM so that they could reasonably be considered in the off medication state once the overnight recording started around 10 PM.

### Memory consolidation task

We used a computerized verbal paired-associates learning task adapted from previous studies investigating sleep-dependent memory consolidation ^32–34^. The task consisted of a learning and a practice block, followed by a pre-sleep and a post-sleep memory recall test (**Figure 1**). Stimuli consisted of 50 pairs of semantically related words (e.g. friend-loyalty; painting-gallery). A French word list was created by translating the English list and was reviewed by a bilingual (native French and English) speaker to qualitatively ensure that no major differences existed in the perceived frequency and difficulty of the words. In the learning block, each word pair was presented on screen for 4 seconds in a random order. In the practice block, the first word of each pair was presented on screen and participants had 3.5 seconds to verbally recall its associate. Following a tone that announced the end of the recall period the correct answer was presented on the screen for 4 seconds. The practice block was immediately followed by a pre-sleep memory recall test where memory for only one half of the word pairs (25 randomly selected pairs) was tested in the same manner, except that the correct answer was not displayed. To prevent active rehearsing, participants were given simple arithmetic problems to solve for 1 minute between blocks. All testing in the evening occurred on average 3:59 hours before sleep. Memory for the remaining half of the word pairs was tested the following morning around 7am. Participants were tested in their first language, either French (n=22) or English (n=13). As age, disease duration, MoCA and memory recall performance did not differ between the two language groups, we conducted all analyses on the full sample. We measured memory consolidation by calculating the overnight relative change in memory performance (i.e., (morning performance – night performance)/night performance).

**Figure 1.**
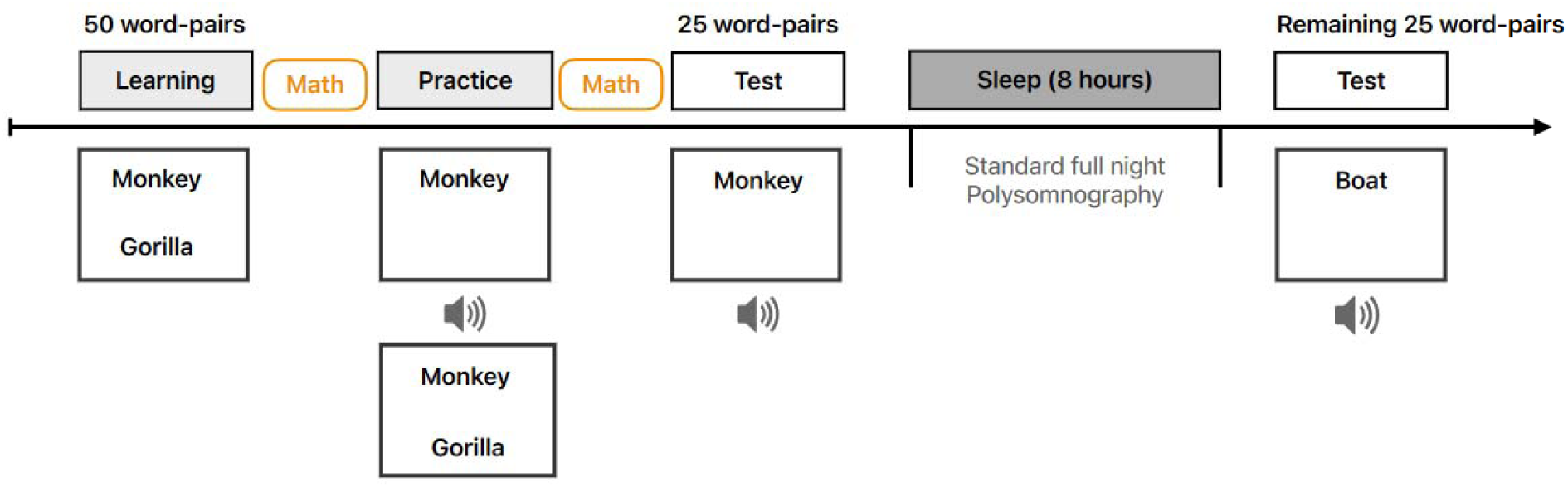
Schematic of experimental design. (**A**) Participants learned 50 semantically related word-pairs in the evening. During the initial learning block, all 50 word-pairs were presented one at a time and participants were instructed to remember them. This was followed by a practice block where one word was presented and participants had to verbally recall its associate. After a tone indicated the end of the recall period, the correct word was presented on screen. The practice block was immediately followed by the pre-sleep recall test where memory for one half of the pairs (randomly selected) was tested. Participants completed simple arithmetic problems for 1 minute between each block. The post-sleep recall test took place the following morning and memory for the remaining 25 pairs was tested. Overnight polysomnography included 6 EEG leads (F3, F4, C3, C4, O1, O2) referenced to the contralateral mastoid, bilateral electrooculogram, submental electromyography.

### Overnight polysomnography

The polysomnography montage consisted of 6 EEG leads (F3, F4, C3, C4, O1, O2), following the 10-20 system and referenced to the contralateral mastoid, a bilateral electrooculogram (EOG), and submental electromyography (EMG). Respiratory inductance plethysmography was used to determine thoracoabdominal motion, and a nasal pressure and oronasal thermistor were used to determine airflow. Signals were recorded using the Polysmith software (Nihon Kohden, Irvin, USA), with a sampling frequency of 200Hz. Sleep stages were visually scored on 30-second epochs following the American Academy of Sleep Medicine (AASM) criteria ^35^. The recording night (i.e., lights were turned off and patients were ready to sleep) started on average at 10:23 PM (SD = ±34 minutes) and finished at 6:00 AM (SD= ±25 minutes). PSG outcomes of interest included total sleep time, sleep efficiency, wake-after sleep onset (WASO) and percentage of time spent in each sleep stage. The scoring of sleep apneas and hypopneas was done according to standard AASM criteria ^36^. The apnea-hypopnea index was measured as the average number of apneas or hypopneas within an hour of sleep.

### Detection of sleep spindles and trains of spindles

The detection of sleep spindles was performed on all EEG leads (F3-M2, F4-M1, C3-M2, C4-M1, O1-M2, O2-M1) during NREM epochs free of artefacts. Artefacts were first detected using an automatic algorithm and, following visual inspection, were rejected from analyses. A band pass filter of 11.1 Hz to 14.9 Hz was applied using a linear phase finite impulse response (FIR) filter (3dB at 11.1 and 14.9 Hz) and the signal was forward and reverse filtered to ensure zero phase distortion and to double the filter order. The root mean square (RMS) of the resulting filtered signal was calculated using a 0.25 second time window and thresholded at the 95^th^ percentile. If at least two consecutive RMS points surpassed the duration threshold of 0.5 seconds, a spindle was detected. Our main variables of interest were sleep spindle density (count per minute), amplitude (µV) and frequency (Hz), as these measures have previously been associated with cognition in patients with Parkinson’s disease ^9,10^.

Spindle trains were counted at each lead on artefact-free periods of sleep and consisted of a minimum of two sleep spindles occurring within 6 seconds ^25^. As we found no significant differences between spindle density or spindle trains between the hemispheres, we averaged the results of the left and right leads together. For spindle trains, we computed three measures: i) proportion of spindles occurring as part of a train, ii) density of spindles occurring as part of a train (i.e., number of spindles part of a train/minutes in sleep stage of interest), and iii) density of spindles occurring outside a train.

### Statistical analyses

To measure the relationship between spindle characteristics and memory consolidation, we computed Pearson’s correlations between each spindle characteristic and overnight change in memory and did this separately for frontal and central leads and for NREM-2 and NREM-3. We present the p-values for the significant associations between spindles and memory consolidation, adjusted for four comparisons (i.e., Leads [Frontal, Central] x Sleep stage [NREM-2, NREM-3]) using the Holm-Bonferroni method. Analyses stratified on allegiance to a train (e.g., density of spindles inside trains vs. density of spindles outside trains) were conducted only for frontal leads during NREM-3 based on the findings in the first set of analyses, and thus were adjusted for two comparisons (see Table S1). We additionally calculated bayes factors to provide an estimate of the evidence in favor of the alternative hypothesis over the null hypothesis for our main results using the BayesFactor package in R ^37^. A Bayes factor between 1 and 3 suggests anecdotal evidence for the alternative hypothesis H_1_, factors between 3 and 10 suggest substantial evidence for H_1_ and factors between 10 and 30 suggest strong evidence for H_1_. All analyses were conducted in R version 4.0.5 and RStudio version 1.4.1106.

**Table 1I.**
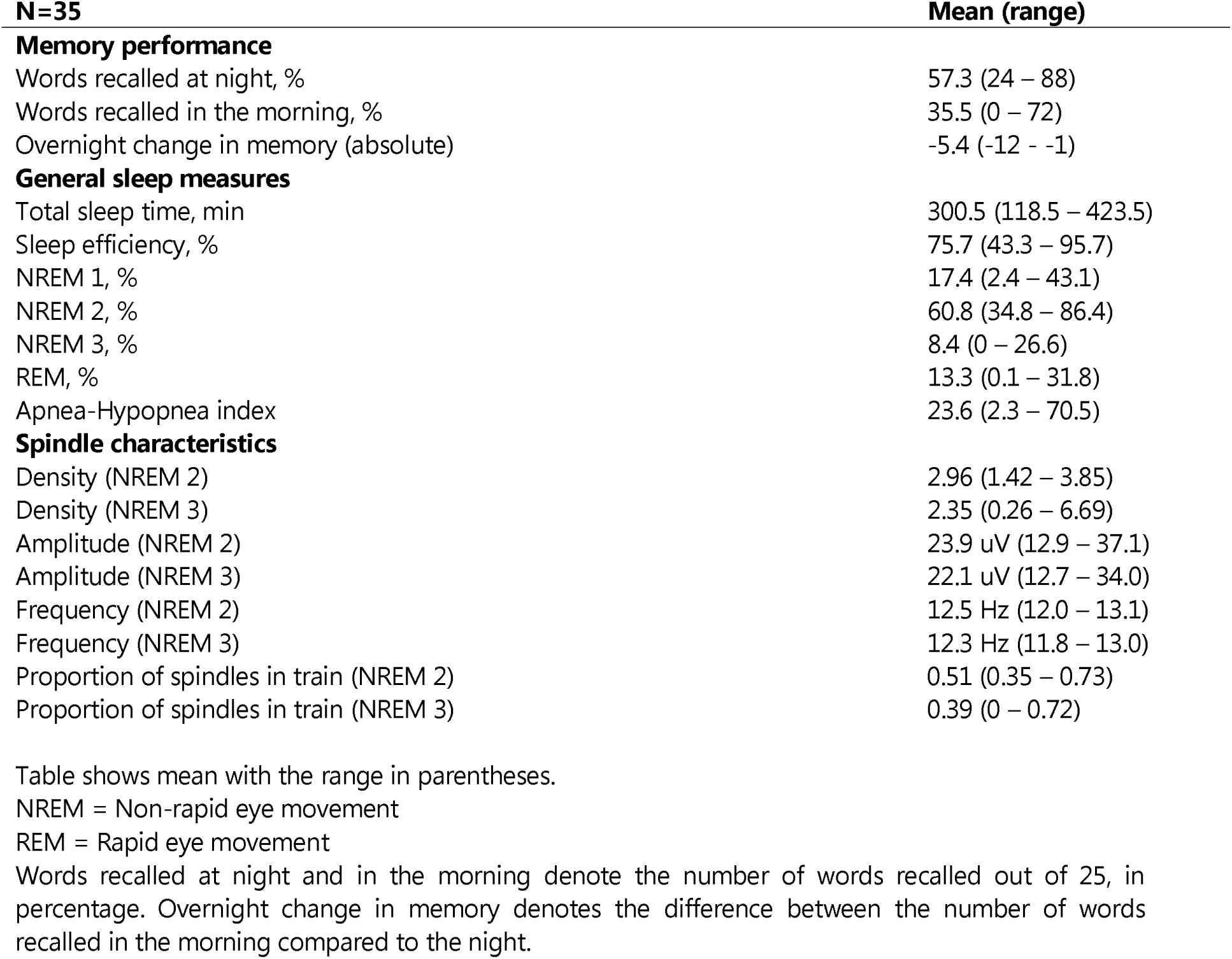
Memory performance and polysomnography.

## Results

### Relationship between spindle density and memory consolidation

The average overnight relative change in memory was −0.41 (SD=0.0023, range −0.0007 to 1) indicating that, as expected, participants correctly recalled fewer words in the morning (mean=8.8 words, SD=4.7) than at night (mean=14.3 words, SD=4.1). We found a significant relationship between spindle density measured in frontal leads during NREM-3 and the overnight relative change in memory such that a higher spindle density was associated with less reduction in memory overnight (R=0.46, p=0.006, p_adj_=0.024, BF_10_=9.8), but not for spindles measured at central leads, nor for spindles measured during NREM-2 (Central NREM2: R=−0.04, p=0.82, BF_10_=0.81; Central NREM-3; R=0.2, p=0.27, BF_10_= 0.64; **Figure 2** and **Supplementary Table 1**). Because the specificity of the effect to NREM-3 sleep and to the frontal leads was somewhat unexpected we ran additional analyses to examine whether significant differences between frontal and central leads in NREM-2 and NREM-3 existed (such as total spindle count, spindle density, artefact-free periods) that could potentially explain this specificity. Overall, no significant differences were found. These analyses are presented in the Supplement and in **Supplementary Figure S1**.

**Figure 2.**
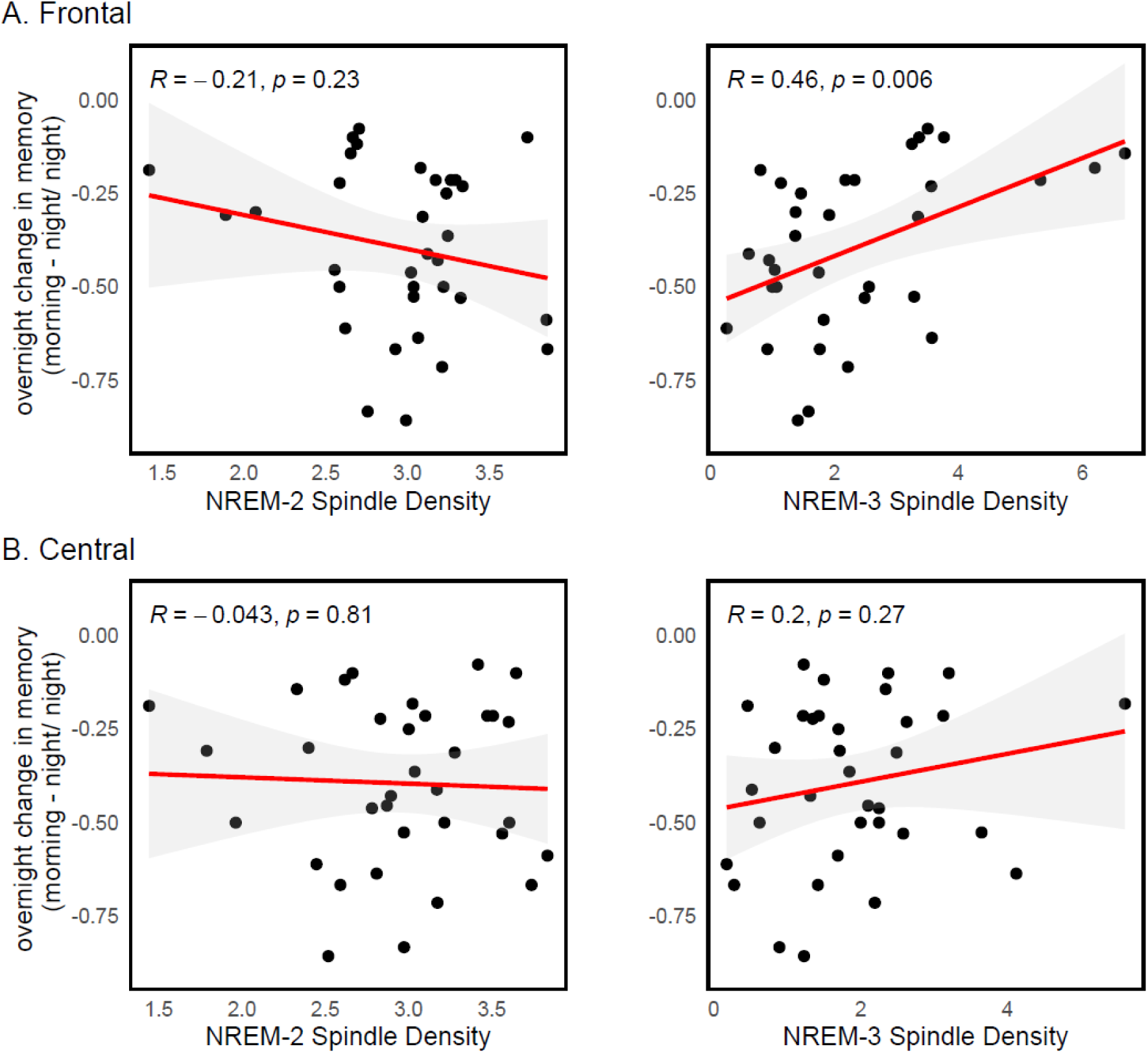
Relationship between sleep spindle density and memory consolidation. **Spindle density is shown for** A) frontal and B) central leads during both NREM-2 and NREM-3 sleep in association with the overnight relative change in memory, where negative values represent worse memory performance in the morning than at night. Higher density (# spindles/min of sleep) of frontal sleep spindles in NREM-3 was associated with less overnight forgetting (R=0.46, p=0.006). Red line denotes the regression line, and shaded areas represent the 95% confidence interval.

### Relationship between spindle trains and memory consolidation

Because of prior evidence suggesting a role for the temporal clustering of spindles in memory consolidation ^25,27,28^, we examined the relationship between spindle trains and overnight memory change. We found that a higher proportion of spindles occurring in trains (as opposed to outside of trains) in NREM-3 at frontal leads was associated with less overnight reduction in memory performance (R=0.46, p=0.0057, p_adj_=0.02, BF_10_=10.13; Figure 4C). This effect was again specific to frontal spindles occurring in NREM-3 sleep. There were no statistically significant associations between spindle trains (neither number of trains nor proportion of spindles in trains) during NREM-2 and the overnight change in memory (Number of trains: R=−0.19, p=0.26; Proportion of spindles in trains: R=−0.23, p=0.17; BF ranging from 0.47 to 0.65; see Table S1). In addition, we separately computed spindle density for spindles occurring as part of a spindle train and for spindles occurring independent of a train. We found that higher density of spindles occurring *inside* a train was associated with less overnight reduction in memory performance (R=0.52, p=0.002, p_adj_=0.004, BF_10_=27.9; Figure 4A), whereas there was no statistically significant relationship between density of spindles occurring *outside* of trains and overnight memory (R=0.2, p=0.2, BF_10_=0.7; Figure 4B).

We also measured the absolute number of trains that occurred in NREM-2 and NREM-3, but the results did not show any significant association with overnight memory change (Frontal leads in NREM-3: R=−0.05, p=0.8, BF_10_=0.4; Figure 4D. Frontal leads in NREM-2: R=−0.16, p=0.39, BF10=0.5; see Table S1). Finally, we compared the characteristics of spindles occurring inside trains versus those occurring outside trains and found no difference with regards to the amplitude and frequency of the spindles (amplitude: t=−0.09, p=0.92; frequency: t=−0.76, p=0.44). However, we found a trend for the duration of the spindles, such that spindles inside trains were longer than spindles outside trains (duration: t=−1.85, p=0.07). Overall, these results suggest that the clustering of spindles into trains is important for the overnight consolidation of declarative memory and this effect does not seem to depend on differences in the characteristics of the spindles.

**Figure 3.**
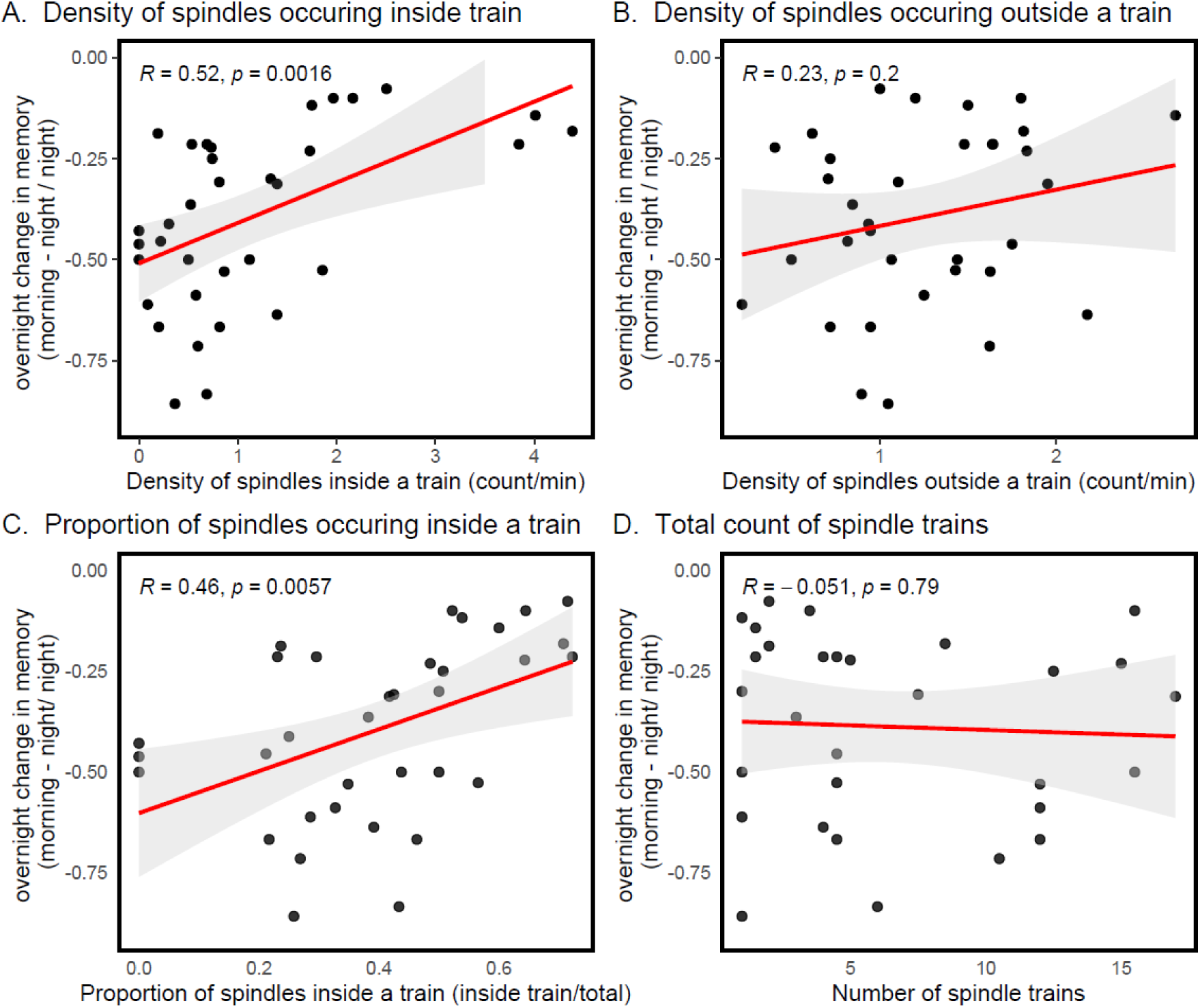
Relationship between memory consolidation and clustering of spindles in NREM-3 at fontal leads. **(A)** Relationship between density of spindles occurring inside trains (count/min) and change in memory overnight, where negative values indicate worse performance in the morning than at night. Increased density of spindles occurring inside trains was associated with less reduction in memory overnight (R=0.52, p = 0.001) **(B)** Density of spindles occurring outside of trains was not associated with the overnight change in memory. **(C)** Relationship between proportion of spindles occurring inside trains (relative to the total number of spindles) and change in memory overnight. A higher proportion of spindles occurring inside trains was associated with less reduction in memory overnight (R=0.46, p=0.005). **(D)** Total train count was not associated with change in memory overnight. Red lines denote regression lines and shaded areas denote the 95% confidence intervals.

### Relationship between general measures of healthy sleep and memory consolidation

To determine if the relationships we found between spindle density and spindle trains and memory consolidation were specific to spindles or merely a reflection of overall quality of sleep, we also computed standard measures of sleep architecture and related them to memory consolidation. As expected, we did not find statistically significant associations between memory consolidation and percentage of time spent in each sleep stage (N1%: R=0.17, p=0.33; N2%: R=−0.04, p=0.79; N3%: R=−0.16, p=0.36; REM%: R=0.04, p=0.81), number of minutes spent in each sleep stage (N1: R=0.02, p=0.87, N2: R=−0.05, p=0.74, N3: R=−0.18, p=0.29, REM: R=0.008, p=0.96), total sleep time (R=−0.06, p=0.74), and sleep efficiency (R=0.03, p=0.86). Because obstructive sleep apnea is common in Parkinson’s disease and is associated with cognitive performance ^7^, we also computed two standard metrics of obstructive sleep apnea. Neither the apnea-hypopnea index (R=−0.097, p=0.59), which reflects the average number of apneas and hypopneas per hour of sleep, nor the time spent under 90% oxygen saturation (R=−0.27, p=0.18) were associated with overnight memory change.

### Relationship between spindle measures and overall cognitive performance

We also wanted to determine if the relationships we found between spindle density and spindle trains and overnight change in memory were specific to memory consolidation (a sleep-dependent cognitive process) or whether these same measures of spindles were also correlated with overall level of cognitive function, as measured by the MoCA. We found that higher frontal spindle density during NREM-3 was associated with higher MoCA scores (R=0.44, p=0.01, BF_10_=6.5). However, there was not a statistically significant association between the proportion of spindles occurring inside trains and MoCA score (R=0.23, p=0.17, BF_10_=0.8). Unsurprisingly, higher MoCA scores were associated with less overnight reduction in memory performance (R=0.46, p=0.005, BF_10_=10.12).

## Discussion

Changes in sleep micro-architecture are commonly reported in patients with Parkinson’s disease and have been found to be predictive of cognitive decline ^9–11^. However, whether changes in sleep micro-architecture could contribute to cognitive dysfunction by interfering with sleep-dependent cognitive processes or whether they are merely a marker of more severe disease is unclear. In a group of patients with Parkinson’s disease with average disease duration of 3.5 years we measured overnight declarative memory consolidation, a cognitive process known to depend on sleep ^38^. We recorded overnight sleep EEG with a focus on sleep spindles because these are sleep oscillations known to play a role in memory consolidation ^39^. We found that lower spindle density during NREM-3 and lower density and proportion of spindles occurring inside trains of temporally clustered spindles were associated with worse overnight declarative memory consolidation. In contrast, standard measures of sleep macro-architecture (i.e., general measures of sleep health) were not associated with memory consolidation. These findings suggest that reduced spindles, known to be a feature of sleep in Parkinson’s patients, and in particular, reduced clustering of spindles into trains, may play a role in the cognitive deficits observed in Parkinson’s patients, and thus advance our understanding of the importance of preserved sleep architecture for cognition in Parkinson’s disease.

Our results are broadly consistent with findings in healthy younger and older adults showing that higher spindle density is associated with better overnight preservation of memory ^13,15–18,40^. We additionally found that greater clustering of spindles into trains was associated with better consolidation in Parkinson’s patients, which is also consistent with recent work in young and older adults ^25,27,28,41^. One proposed mechanism for the relationship between spindles and memory consolidation is that spindles could facilitate the reactivation of a memory engram and the integration of a memory trace into relevant brain networks during sleep ^14,20,42–45^. Clustered spindles have been proposed to be more efficient than isolated spindles as they might reflect enhanced functional connectivity in cortical-subcortical networks involved in learning ^46^. In addition to the potential importance of the temporal organization of spindles for information processing, some evidence suggests that spindles occurring inside trains could have certain characteristics, such as higher amplitude and longer duration, that reflect a process beneficial for memory consolidation ^26,28^. In our sample, however, the amplitude and duration of spindles occurring inside trains did not differ significantly from those of spindles occurring outside trains. This suggests that, even in the absence of higher amplitudes and longer duration, the temporal organization of spindles may play a role in the consolidation of a memory trace. Overall, by showing that not only a higher density of spindles, but more specifically, a higher density of temporally clustered spindles, are associated with better overnight memory consolidation, our results provide the first evidence of a possible mechanism directly linking sleep alterations to mnemonic cognitive function in Parkinson’s patients.

Our study did not include a sample of healthy older adults for comparison, and therefore our results cannot speak to the specificity of these effects in Parkinson’s patients. Indeed, aging has also been associated with a reduction in spindle density and this reduction has been associated with impaired memory consolidation ^47–49^. However, in Parkinson’s disease, the reduction in sleep spindle density is more pronounced than in healthy older adults, hence suggesting that our findings are particularly relevant to Parkinson’s patients ^10–12,50^. Reduced spindle density in Parkinson’s patients has also been shown to be predictive of future cognitive decline, further highlighting the possible importance of considering the role of altered sleep spindles in the development of cognitive dysfunction in PD. Much less is known about aging effects on the temporal clustering of spindles. One study found that the proportion of clustered spindles and the length of train of clustered spindles decreased with age, and that shorter trains were associated with worse declarative memory consolidation ^28^. Our study is the first to report on sleep spindle trains in Parkinson’s disease, therefore it remains unknown if the effect of Parkinson’s disease on spindle trains is similar to that of the disease on spindle density, i.e. an enhancement of age-related effects. Interestingly, animal studies have shown that noradrenergic activity from the locus coeruleus, a structure known to be affected by neurodegeneration early in Parkinson’s disease ^51^, modulates the clustering of spindles into trains ^52–55^, suggesting that this clustering might be disproportionately affected by Parkinson’s disease. Though future work comparing Parkinson’s patients to older adults is necessary, taken together, our results raise the possibility that the relationship between spindle trains and memory consolidation might be especially relevant to consider in Parkinson’s disease.

We found that higher spindle density was associated with better consolidation during NREM-2 but not NREM-2. Previous findings examining the relationship between spindles and memory consolidation have been inconsistent on this point: a relationship has been found for spindles in NREM-2 ^13,56^ as well as NREM-3 ^57,58^, but most studies seem to collapse across NREM-2 and NREM-3 ^16,22,48,59,60^, hence leaving it unclear whether the associations between spindle density and memory are specific to a particular stage of sleep. It is also unknown whether the functional role of spindles differs in NREM-2 and NREM-3, and whether this has implications for memory consolidation. One possibility is that different types of memory are preferentially consolidated during different stages of sleep. In keeping with our findings, declarative memory consolidation, as measured using tasks similar to ours, has primarily been reported in association with NREM-3 oscillations, and in particular, in association with the coupling that occurs between spindles and slow waves ^24,61,62^. In Parkinson’s disease, most research has focused on deficits in procedural memory consolidation, but even these deficits have not been measured in association with alterations in sleep micro-architecture ^63–65^. Whether our findings linking sleep alterations to declarative memory consolidation would translate to the domain of procedural memory consolidation is an important open question given the relevance of motor learning to the disease and given the potentially modifiable nature of these sleep alterations.

We also found that the relationship of spindle density and spindle trains to overnight declarative memory consolidation was specific to spindles measured at the frontal leads, which is consistent with previous studies in younger adults ^59,60,66^. Two studies that measured both fast (13 – 15 Hz) and slow (11-13Hz) spindles found that the relationship to memory consolidation was specific to slow spindles ^59,60^. In our sample, the majority of spindles detected (at both central and frontal leads) would be classified as slow spindles (68%, refer to **Table S2**). Although evidence for distinct functional roles for fast and slow spindles is mixed ^28,67,68^, the fact that the majority of spindles in our sample were slow spindles could explain why the association we observed with declarative memory consolidation was specific to spindles measures at frontal leads, because spindles measured at this location are predominantly slow spindles ^69^.

We also found that better overall cognitive function, as measured with the MoCA, was associated with increased frontal sleep spindle density during NREM-3, and that higher MoCA scores were associated with better overnight memory consolidation. Because no previous research has examined the relationship between overnight memory consolidation and more general cognitive function (neither in younger nor in older adults) it is unknown how memory consolidation contributes to more general cognitive function, and whether sleep supports both separately or whether sleep’s beneficial effect on consolidation supports more general cognitive function by, for instance, reducing the cognitive resources required to re-learn information. Our study was not designed to address this because our only measure of general cognitive function was the MoCA, which is designed to be a screening tool for cognitive impairment and is not designed to provide a continuous linear measure of cognitive function ^70^. Nonetheless, the fact that performance on the MoCA, which combines multiple domains of cognitive function and includes only a test of short-term memory, was associated with the overnight *change* in memory normalized to baseline (short-term) memory performance is interesting and will require further study.

There were several limitations to our study that are important to note. First, our sample of Parkinson’s patients included patients with and without obstructive sleep apnea, a sleep disorder that is very common in Parkinson’s disease, occurring in up to 60% of patients. In our sample, 65% of participants were considered to have moderate to severe obstructive sleep apnea. Obstructive sleep apnea has been associated with poor cognitive function ^7,71,72^ and it is possible that it could also affect sleep oscillations ^29^. For instance, obstructive sleep apnea is known to fragment sleep ^73^, which in turn could affect sleep spindles and other sleep oscillations by reducing the amount of uninterrupted sleep. Second, our relatively small sample size prevented us from having the statistical power to address the clinical heterogeneity in Parkinson’s disease, and to conduct stratified analyses to examine the contributions of different clinical characteristics such as obstructive sleep apnea, disease severity and baseline cognitive function, all factors that might be expected to influence the relationship between sleep micro-architecture and memory consolidation.

In summary, we found that lower sleep spindle density, already known to be reduced in Parkinson’s disease to a greater extent than in healthy older adults, and lesser spindle clustering were associated with worse overnight declarative memory consolidation, a specific sleep-dependent cognitive process. This represents an important first step towards delineating the effects of Parkinson’s disease-related sleep impairments on cognition in Parkinson’s patients. Our results lend support to the recently observed relationship between abnormal sleep spindles and the risk of developing cognitive impairment, but several questions remain unanswered. First, future work will be required to determine if impaired memory consolidation in itself could represent a mechanism underlying cognitive impairment, by resulting, for instance, in the diversion of cognitive resources towards the repeated encoding of information that would otherwise already be consolidated and remembered. Alternatively, memory consolidation may be but one of many cognitive processes directly affected by reduced spindle density and clustering, all of which could contribute to more global cognitive impairment. These questions are all the more important considering that impaired sleep micro-architecture is highly prevalent in Parkinson’s disease and potentially modifiable through both existing pharmacological interventions, and through more recently described non-pharmacological interventions such as closed-loop auditory stimulation during sleep ^74–76^.

## Acknowledgements

We thank the participants for their time and commitment, Marianne Gingras and Sena Tavukçu for their assistance with data collection, and the Quebec Parkinson Network for assistance with participant recruitment.

## Funding

This work was supported by a Parkinson Canada grant (MS), Weston Family Foundation grant (MK) and by the Fonds de recherche du Québec – Santé (MS).

## Competing interests

The authors declare no competing interests.

## Authorship contributions

**Soraya Lahlou:** Formal analysis, Writing – Original Draft. **Marta Kaminska:** Funding Acquisition, Writing – Review & Editing; **Julien Doyon:** Methodology, Writing – Review & Editing; **Julie Carrier:** Methodology, Supervision, Writing – Review & Editing; **Madeleine Sharp:** Conceptualization, Methodology, Resources, Writing – Review & Editing, Supervision, Project Administration, Funding Acquisition.

## Notes

### Competing Interest Statement

The authors have declared no competing interest.

