## Supplemental Material for "Sleep spindle density and temporal clustering are associated with sleep-dependent memory consolidation in Parkinson’s disease"

**Supplementary Material**

|  | **NREM-2** | **NREM-3** |
| --- | --- | --- |
| Spindle density (Frontal) | R= -0.22, p=0.20, BF_10_=0.76 | **R=0.46, p=0.006, p_adj_=0.02, BF**_10_**=9.78** |
| Spindle density (Central) | R= -0.04, p=0.82, BF_10_=0.81 | R=0.2, p=0.27, BF_10_=0.64 |
| # Spindle train (Frontal) | R=-0.19, p=0.26, BF_10_=0.65 | R= -0.05, p=0.79, BF_10_=0.41 |
| # Spindle train (Central) | R=-0.12, p=0.46, BF_10_= 0.47 | R=-0.16, p=0.39, BF_10_=0.55 |
| Proportion of spindles in trains (Frontal) | R=-0.23, p=0.17, BF_10_=0.84 | **R=0.46, p=0.0057, p_adj_= 0.02, BF**_10_**=10.13** |
| Proportion of spindles in trains (Central) | R=-0.13, p=0.42, BF_10_=0.49 | R=0.17, p=0.31, BF_10_=0.58 |
| Spindle density (# spindles in trains/min) | Not measured | **R=0.52, p=0.002, p_adj_=0.004, BF_10_=27.9** |
| Spindle density (# spindles outside trains/min) | Not measured | R=0.23, p=0.2, BF_10_=0.77 |

|  | **NREM-2** | | **NREM-3** | |
| --- | --- | --- | --- | --- |
|  | **Slow** | **Fast** | **Slow** | **Fast** |
| Frontal spindles (n ±SD) | 196 ±21.2 | 47 ±36.4 | 21.2 ±19.9 | 3.67 ±2.84 |
| Central spindles (n ±SD) | 138 ±71.3 | 107 ±80.9) | 13.7 ±15.5 | 9.04 ±8.62 |
| Frontal spindles (% ±SD) | 79.8% ±13.1 | 20.2% ±13.1 | 82.0% ±11.9 | 18.0% ±11.9 |
| Central spindles (% ±SD) | 57.4% ±19.2 | 42.6% ±19.2 | 55.1% ±20.9 | 44.9% ±20.9 |

**Table S1.** Correlations between sleep spindles and memory consolidation. P_adj_ denotes p-values adjusted for 4 comparisons (except for spindle density inside and outside trains) using the Holm-Bonferroni method.

**Table S2.** Number and percentage of slow and fast sleep spindles across frontal and central leads for NREM sleep. Sleep spindles were detected in the range of 11.1 and 14.9 Hz. Then, spindles with a frequency of ≤12.99 Hz were categorized as slow sleep spindles, and spindles with a frequency of ≥13.00 Hz were categorized as fast sleep spindles

**Exploratory analyses on central and frontal leads during NREM-2 and NREM-3**

We compared spindle density and total count of spindles in both frontal and central leads and during both NREM-2 and NREM-3 (Figure 3), and compared the minutes of artefact-free periods, which is the amount of time devoid of EEG artefact on which the spindle detection was performed. These analyses were conducted to rule out the possibility that the specificity of the results to frontal leads is not due to relatively lower noise at these leads. Interestingly, we found that, although there are no significant differences in total spindle count between frontal and central leads in NREM-3 (paired t-test: t=-1.55, p=0.13), we found that there is a significantly higher spindle density at frontal leads compared to central leads in NREM-3 (paired t-test: t=-2.53, p=0.01), though this does not survive correcting for multiple comparisons.

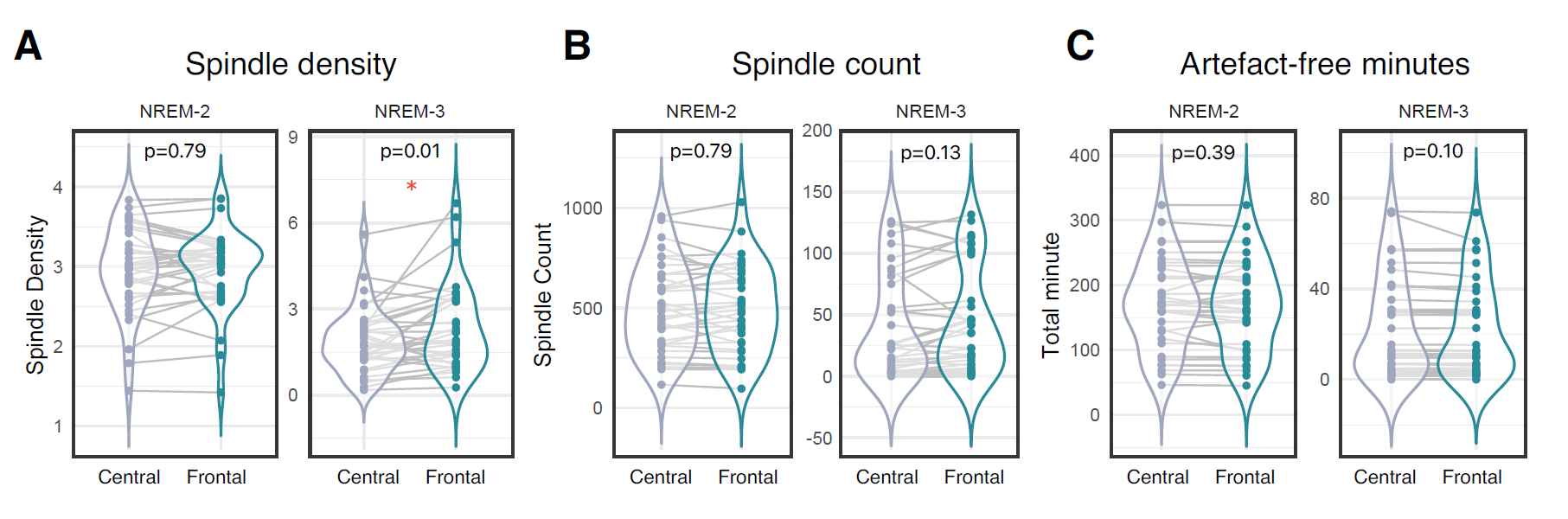

**Figure S1.** **Comparing sleep characteristic between central and frontal leads during NREM-2 and NREM-3.** **(A)** Spindle density, **(B)** count, and **(C)** total artefact-free minutes measured at central and frontal leads during NREM-2 and NREM-3 stages. Artefact-free minutes reflect the minutes free of EEG artefacts where the spindle detection was conducted. **(A)** Spindle density during NREM-3 was higher at frontal leads than at central leads (paired t=-2.61, p=0.01). There was no difference in spindle density between central and frontal leads during NREM-2 (paired t=-0.26, p=0.79). **(B)** No differences in total spindle count between central and frontal leads during NREM-2 (paired t=0.26, p=0.79) and NREM-3 (paired t=-1.57, p=0.13). **(C)** No differences in artefact-free minutes of sleep between central and frontal leads during NREM-2 (paired t=0.86, p=0.40) and NREM-3 (paired t=1.66, p=0.10). Grey lines represent a single participant.
